# Human Tau Isoform Aggregation and Selective Detection of Misfolded Tau from Post-Mortem Alzheimer’s Disease Brains

**DOI:** 10.1101/2019.12.31.876946

**Authors:** Ling Wu, Zerui Wang, Shradha Ladd, Darren T. Dougharty, Sidharth S. Madhavan, Madeleine Marcus, Frances Henderson, W. Keith Ray, Christopher Tan, Sandra Siedlak, Jianyong Li, Richard F. Helm, Xiongwei Zhu, George S. Bloom, Wen-Quan Zou, Bin Xu

## Abstract

Tau aggregates are present in a large number of neurodegenerative diseases known as “tauopathies”, including Alzheimer’s disease (AD). As there are six human tau isoforms in brain tissues and both 3R and 4R isoforms have been observed in the neuronal inclusions, we tested whether tau isoforms behave differently in aggregation. We discovered that all six tau isoforms are capable of forming PHF-tau like filaments and the 3R tau isoforms aggregate significantly faster than their 4R counterparts. We further mapped key segments of tau isoforms that contribute to their aggregation kinetics, where it was determined that microtubule binding domains R2 and R3 were the major contributors to tau aggregation. To evaluate the feasibility of using the six recombinant tau isoforms as substrates to amplify misfolded tau, we demonstrated that full-length human tau isoforms can seed and detect misfolded tau from the post-mortem AD brain tissues with high specificity by an ultrasensitive technology termed real-time quaking-induced conversion (RT-QuIC). Mass spectrometric analysis of PHF-tau samples extracted from AD brains identified peptides corresponding to all major forms of human brain tau isoforms along with a consensus hyperphosphorylated peptide near the C-terminus. Together, our findings not only reveal new aggregation kinetic properties of human tau isoforms, support the development of methods to quantitatively measure misfolded human tau isoforms in AD brains, but also uncover the capability of full-length human tau isoforms as substrates for “prion-like” tau seeding by RT-QuIC assays that may be used for new biomarker development for AD and other tauopathy diagnosis.

## Introduction

Alzheimer’s disease (AD) is now the most expensive disease in the US, exceeding even cancer and cardiovascular disease for the total annual cost to American society (Hurd et al, 2013). AD is characterized by the accumulation in the brain of two types of abnormal structures in the brain, extracellular Aβ amyloid plaques and intraneuronal tau neurofibrillary tangles (Selkoe et al, 2001). Until recently, plaques and tangles were thought not only to represent molecular hallmarks of AD, but also to cause the synaptic dysfunction and neuronal death that lead to the memory and cognitive deficits characteristic of AD patients (Bloom, 2014; Spires-Jones & Hyman, 2014). Several lines of evidence have suggested that pathological changes in tangles correlate better with neuronal dysfunction than Aβ deposits (Wilcock & Esiri, 2012; Nelson et al, 2012). Moreover, a close relationship between tau aggregates and neuronal loss is well established in hippocampus and cerebral cortex tissues (Goedert & Spilantini, 2017).

Tau aggregates are present in a large number of neurodegenerative diseases known as “tauopathies” (Ghetti et al, 2015). They include AD, Pick’s disease (PiD), progressive supranuclear palsy (PSP), corticobasal degeneration (CBD), as well as frontotemporal dementia and Parkinsonism linked to chromosome 17 (FTDP-17), the latter being caused by mutations in *MAPT* (tau). Six tau isoforms are expressed in adult human brain, produced by alternative mRNA splicing of transcripts from *MAPT* gene. These isoforms contain either three or four microtubule-binding repeats (3R or 4R tau) and 0-2 N-terminal inserts (0N, 1N, or 2N tau) (Goedert et al, 1989; Fig. 1A). Significantly, isoform composition and morphology of tau filaments can differ between tauopathies, suggesting the existence of distinct tauopathy strains. In AD, both 3R and 4R isoforms make up the neuronal inclusions, whereas in PiD, 3R isoforms predominate in the neuronal deposits. The assembly of 4R tau into filaments is a characteristic of PSP and CBD. Transmission of the AD pathology and related tauopathies is believed to be through the “prion-like” mechanisms that ultimately yield intercellular spreading of protein aggregates. Aggregate propagation requires protein release into the extracellular space, uptake by connected cells and seeded aggregation of soluble cellular proteins. Currently, the mechanisms underlying the prion-like seeding and spreading of different tau isoforms are not fully understood.

**Figure 1.**
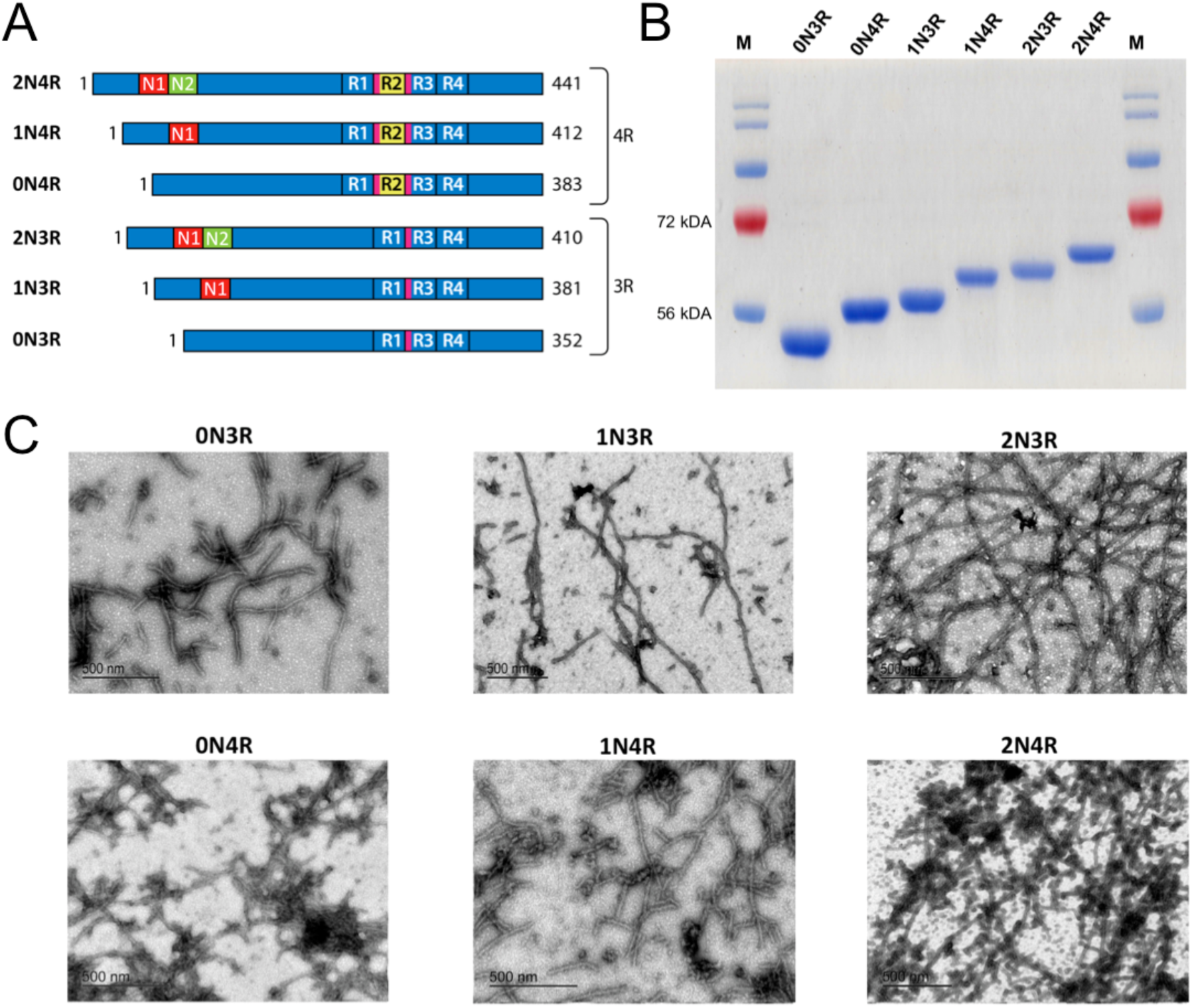
Human tau isoforms. (A) Schematic illustration of the alternative spliced six isoforms of human CNS tau. Amino-terminal insert domains (N1 and N2) and carboxy-terminal repeat domains (R1-R4) are shown. Amino acid residue numbers for each isoform are listed. Locations for the hexapeptide sequences (VQIINK in R2 and VQIVYK in R3) are highlighted in magenta. (B) SDS-PAGE analysis demonstrated six recombinant human tau isoforms purified to homogeneity. (C) TEM images of mature filaments of all six human tau isoform are shown. 30 μM of each tau isoform was prepared in 20 mM Tris pH 7.4 with 0.06 mg/ml heparin and incubated at 37 °C for 12 days. Mature filaments were visualized after staining with uranyl acetate. Size bars are marked.

**Figure 2.**
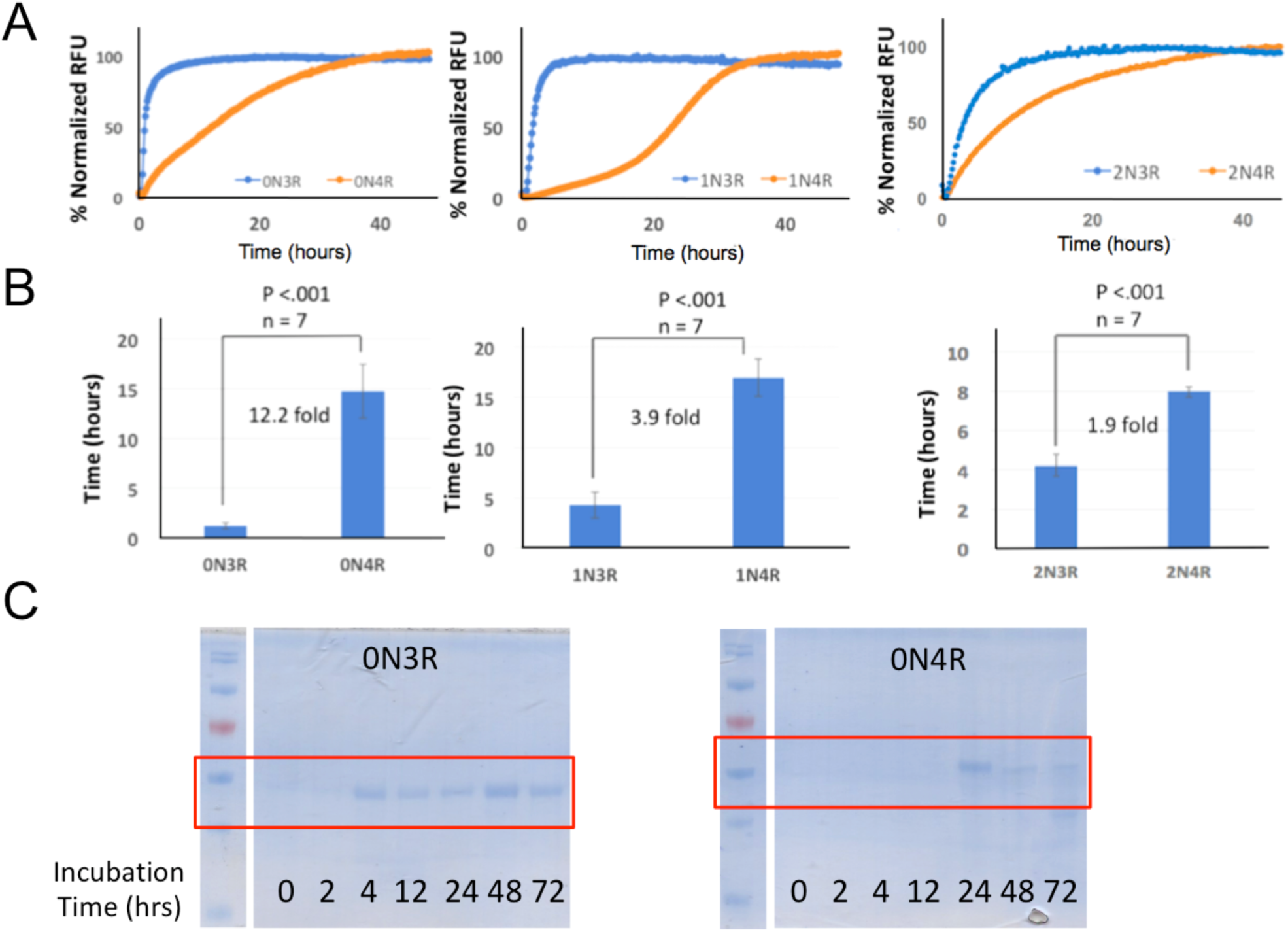
Pair-wise aggregation kinetics comparison of 3R and 4R tau isoforms. (A, B) Quantitative analysis of tau isoform aggregation kinetics using ThT fluorescence-based assays. 30 μM of 0N3R or 0N4R (Left Panel), 1N3R or 1N4R (Center Panel), and 2N3R or 2N4R (Right Panel) were incubated in 20 mM Tris pH 7.4 with 0.06 mg/ml of heparin at 37 °C for specified time periods. Kinetic fluorescence traces are shown in panel A. Corresponding t_1/2_ values for each aggregation kinetics curves are shown in panel B. (C) Aggregation kinetics comparison of 0N3R and 0N4R isoforms by Sarkosyl-insoluble tau pelleting assay. 30 μM of 0N3R or 0N4R were incubated in 20 mM Tris pH 7.4 with 0.06 mg/ml of heparin at 37 °C for specified time periods. Aggregated tau isoforms (Sarkosyl-insoluble tau) with equal loading volumes were visualized by SDS-PAGE after Commassie Blue staining.

Alternative splicing of *MAPT* exon 10 gives rise to tau isoforms with three (3R) or four (4R) microtubule (MT) binding repeats. Normal adult humans are thought to express approximately equal levels of the 3R-tau and 4R-tau isoforms; however, select sporadic tauopathies exhibit imbalanced of 4R-tau isoform deposition within neurofibrillary tangles (NFTs) (Arai et al, 2001; de Silva et al, 2003). Increased 4R-tau expression without change to total tau in hTau mice induced more severe seizures and nesting behavioral abnormalities, increased tau phosphorylation and oligomers (Schoch et al, 2016). In more than 50 disease-causing mutations found in the *MAPT* gene, about one-third of them affect exon 10 splicing and consequently the normal 3R/4R balance (Niblock and Gallo, 2012). The majority of those mutations favor exon 10 inclusion, leading to an increase in 4R (Spillantini and Goedert, 2013), but mutations yielding 3R excess were also linked with disease, suggesting that tau isoform imbalance is detrimental regardless of whether the ratio shifts toward 3R or 4R (Andreadis, 2012). Modulating the 3R/4R ratio in the prefrontal cortex led to a significant reduction of pathological tau accumulation with improvement of neuronal firing and reduction of cognitive impairment in mice (Espindola et al, 2018).

Many protein misfolding diseases, such as AD and other tauopathies (Caughey & Lansbury, 2003), Parkinson’s disease (Lee & Trojanowski. 2006), type 2 diabetes (Velander et al, 2017; Wu et al, 2017), involve analogous pathological accumulation of disease-specific amyloidogenic protein in the form of self-seeding filaments or sub-filamentous deposits. Two methods to amplify misfolded proteins *in vitro*, the real-time quaking-induced conversion (RT-QuIC) assay and the protein misfolding cyclic amplification protocol (PMCA) are used for ultrasensitive detection in a number of neurodegenerative diseases, such as prion disease (Moda et al, 2014; Orru et al, 2017; Wang et al, 2019), Aβ oligomers in AD (Salvadores et al, 2014), and α-synuclein in Parkinson’s disease (Fairfoul et al, 2016; Shahnawaz et al, 2017; Groveman et al, 2018). Specifically for misfolded tau detection in AD and other tauopathies such as Pick’s disease, detection of the prion-like tau seeding activities by RT-QuIC assays was successfully developed using engineered short fragments of tau or a mixture of tau fragments with mutations as a substrate (Saijo et al, 2017; Kraus et al, 2019). However, detection of AD and related tauopathies with recombinant full-length human tau isoforms that match tau isoforms *in vivo* has not been reported.

In this study, we produced all six recombinant human tau isoforms that are as aggregation-competent to form oligomers and mature filaments as those in AD and other tauopathies (Swanson et al, 2017). We dissected the key segments in tau isoforms contributing to their aggregation. Our kinetic work revealed that 3R tau isoforms aggregate significantly faster than their 4R counterparts. Mass spectrometric analyses of PHF-tau samples showed the feasibility for targeted and quantitative measurements of tau isoforms in AD brains. Using the ultrasensitive RT-QuIC assay, we demonstrated prion-like seeding activities of misfolded tau from post-mortem AD brains with both recombinant 3R-tau and 4R-tau isoforms as substrates with high specificity, suggesting the potential uses of various recombinant human tau isoforms for developing RT-QuIC-based diagnosing, characterizing, and predicting consequences of AD and non-AD tauopathies.

## Results

### Recombinant human tau isoforms are capable of forming PHF-tau like filaments

We first established a system to produce all six recombinant human tau isoforms and developed multiple *in vitro* biochemical and biophysical assays to define and monitor tau oligomer/amyloid formation. Expression vectors for his-tagged versions of all six wild-type human tau isoforms (Fig. 1A) were kindly provided by the late Dr. Lester “Skip” Binder and Dr. Nicolas Kanaan of Michigan State University. All tau isoforms and truncation mutants were purified to homogeneity (>90% purity) by Ni-NTA affinity chromatography and additional steps of size exclusion chromatography (Figs. 3B, 3E, and 3H). Full-size tau isoforms went through an additional boiling step for enhanced purity (>95%; Fig. 1B; Borghorn et al, 2005). Recombinant human tau isoforms were capable of generating circular oligomers as well as filaments that can be visualized by transmission electron microscopy (TEM; Fig. 1C). They have similar morphology as those reported in the literature (Patterson et al, 2011) and as those of PHF tau filaments extracted from AD brains (Fig. 6C; Goedert et al, 1992; Guo et al, 2016). We noticed that it takes significantly longer time period (typically 10-12 days under incubation conditions we used) to form fibrillar filaments, in marked contrast to other more amyloidogenic proteins such as amylin, for which it takes only 2-3 days to form fibrils under similar buffer conditions (Velander et al, 2016; Wu et al, 2017). It is possible that different isoforms may take varying length of time in fibrillation. Required long period of incubation in forming mature tau filaments reflects relative stability of the tau isoforms in solution in comparison with other amyloidogenic proteins. This is also consistent with the unusual step of boiling whole cell lysates in purification of recombinant tau isoforms (Barghom et al, 2005).

**Figure 3.**
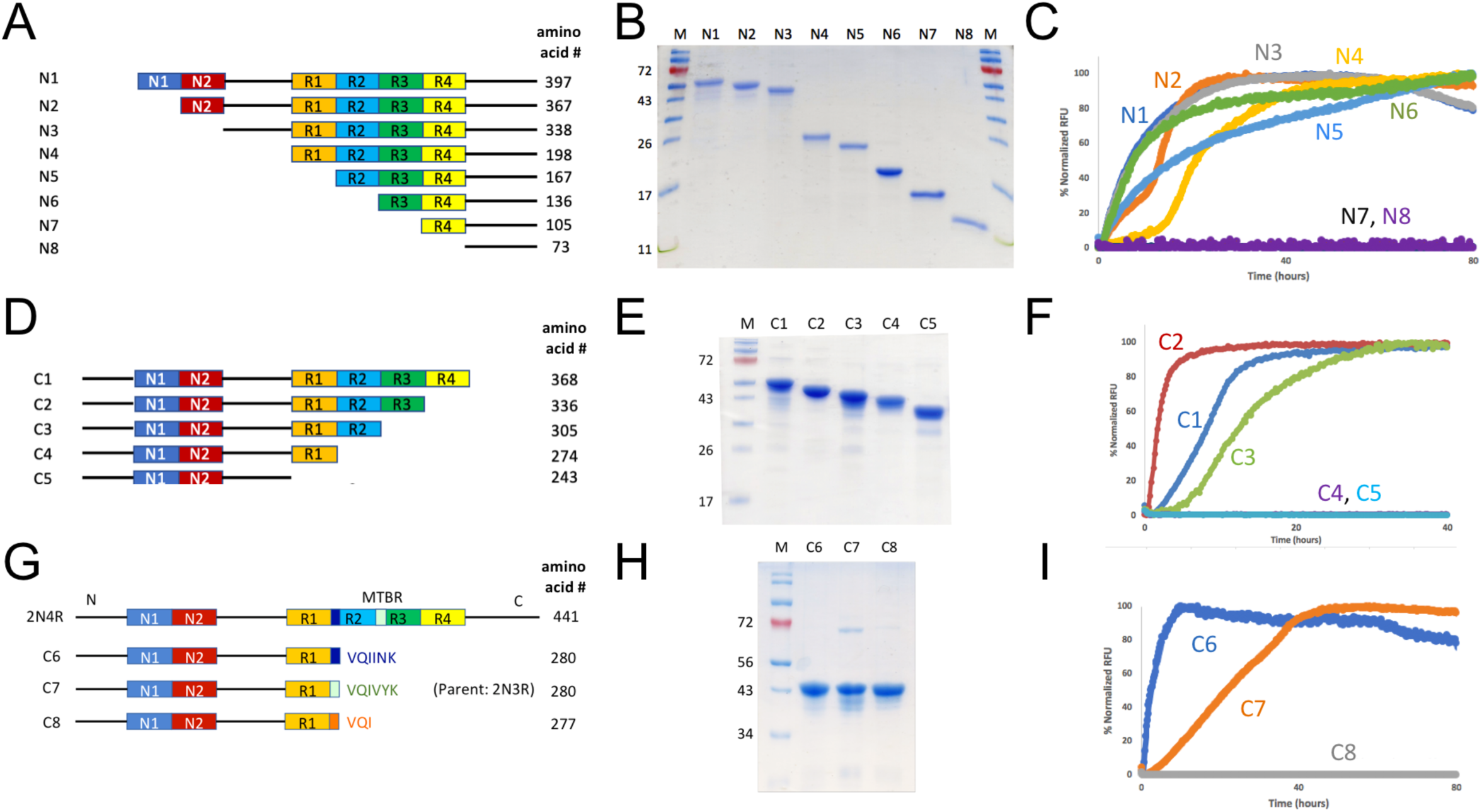
Mapping the key segments in 2N4R tau for aggregation using truncation mutagenesis. (A, D, G) Each construct is schematically illustrated. Except for C7 construct, which was cloned using 2N3R tau as the template, all other constructs were cloned using 2N4R tau as the template. A his_6_-tag was added to each construct in the carboxy-terminal (not shown) to facilitate protein purification. Amino acid numbers for each construct are listed. (B, E, H) SDS-PAGE of purified recombinant tau proteins for corresponding constructs. (C, F, I) ThT fluorescence assays for each truncation mutant. 15-30 μM of tau mutant proteins were incubated in 20 mM Tris pH 7.4 with 0.06 mg/ml of heparin at 37 °C for specified time periods. The results are representative of at least triplicated experiments.

### 4R-tau isoforms have significant delayed aggregation kinetics than each of their 3R-tau counterpart

A ThT fluorescence assay was employed to characterize tau aggregation kinetics. Recombinant human tau variants were aggregation-competent as shown in thioflavin T (ThT) fluorescence assays (Fig. 2A). The ThT assay time course profile provided precise information on aggregation kinetics including t_1/2_ measurements. All the 4Rs variants reached a plateau near or after 40 hours incubation, which were significantly slower than the 3R isoforms (2-20 hours, Fig. 2A). Pair-wise t_1/2_ comparison of 3R and 4R isoforms revealed significant difference in aggregation kinetics: 0N3R and 0N4R have t_1/2_s of 1.2 ± 0.4 h and 14.7 ± 3.0 h respectively, a drastic 12.2-fold change; The 1N3R and 1N4R isoforms have t_1/2_s of 4.3 ± 1.1 h and 17.0 ± 1.6 h respectively, a 3.9-fold change; and 2N3R and 2N4R have t_1/2_s of 4.2 ± 0.6 h and 8.0 ± 0.3 h respectively, a 1.9-fold change (Fig 2B). These results demonstrated that R2 repeat domain bearing 4R tau isoforms form aggregates much slower compared to the R2-lacking 3R isoforms, rate differences that may reflect the sequence variations of 0N, 1N and 2N isoforms, specifically the N1 and N2 domains.

To further verify our findings, we performed Sarkosyl-insoluble tau pelleting assays to assess tau assembly kinetics (Taniguchi et al, 2005; Aoyagi et al, 2007). To compare the nucleation properties of 0N3R and 0N4R tau isoforms, we examined the lag time of fibrillation in the absence or presence of heparin. No Sarkosyl-insoluble tau was observed with either isoform in the absence of heparin (data not shown). Incubation of tau with heparin resulted in an increase of Sarkosyl-insoluble tau in both isoforms (Fig. 2C). Although 0N4R did not exhibit aggregation for up to 24 hours, 0N3R tau started to assemble at 4 hours using identical fibril formation conditions, in good agreement with the results from ThT fluorescence assays. Sarkosyl-insoluble tau pelleting assay was not sensitive enough to detect the difference between 1N3R/1N4R and 2N3R/2N4R pairs, presumably due to their less pronounced kinetic differences than that of 0N3R/0N4R (data not shown).

### Truncation mutagenesis identifies R2 and R3 as key contributors for tau isoform aggregation

To systematically investigate the roles of individual domain of the tau protein in aggregation, a series of truncation mutants based on 2N4R tau template shortened from either the N-terminus and the C-terminus were generated (Figs. 3A and 3D) and purified to homogeneity (Figs. 3B and 3E). We performed ThT fluorescence assays to test each variant for their amyloid forming capability. In all the N-terminal variants, only N7 and N8 variants were not able to generate significant fluorescence, suggesting these two variants were not capable of forming tau amyloid (Fig. 3C). Similarly, C-terminal variants, C4 and C5 were not able to form tau amyloid (Fig. 3F). Taken together, these results indicated that MT binding repeat domains R2 and R3 play key roles in tau isoform aggregation: retaining any one of these two domains (such as variants N6 or C3) render an amyloidogenic mutant. To further pinpoint if the hexameric sequences of “VQIINK” from the R2 domain or “VQIVYK” from the R3 domain are critical sequence)(s) in aggregation, we engineered three constructs, C6, C7 and C8 (Fig. 3G). Three constructs have the same N-terminal sequences up to the R1 repeat domain. The differences are the extended hexapeptide sequences of “VQIINK” (C6), “VQIVYK” (C7), or tripeptide sequence “VQI” (C8). Consistent with literature, inclusion of “VQIINK” or “VQIVYK” sequence renders each amyloidogenicity, but extension of “VQI” sequence is not sufficient (Fig. 3I). Our results not only verified the key roles of these two hexapeptide sequences in tau aggregation (von Bergen et al, 2000; von Bergen et al, 2001; Li & Lee, 2006), but also provided a new insight that inclusion of these hexapeptide sequences is both necessary and sufficient for tau aggregation.

### Full-length tau isoform based RT-QuIC assay selectively detects misfolded tau seeds in the AD brain but not in non-AD brains

RT-QuIC assay has been used for detection of seeding activities of misfolded proteins, first for prions and then for αSyn and tau *in vitro* (Atarashi et al., 2008; Orru et al., 2017; Groveman et al., 2018; Sano et al., 2018; Saijo et al., 2017; Kraus et al., 2019). To determine whether various recombinant wild-type human tau isoforms can be converted by the tau aggregate seeds from AD brain samples, we detected autopsied AD and non-AD brain samples from neuropathologically confirmed cadavers (Table 1). Since there are six tau isoforms present in the human brain (Espinoza et al, 2008), we tested how individual isoforms worked as substrates for the RT-QuIC assay of tau aggregation. The tau seeding activities of brain samples from multiple AD and non-AD patients as seeds in the presence of individual six recombinant tau isoforms are shown in Figure 4 and related statistical analyses are shown in Figure 5. Tau-seeding activities were detected starting at approximately 10-30 hours and reached a plateau at about 60 hours. In contrast, non-AD samples showed no or minimal seeding activities for the entire 60 hours period and the seeding activities were significantly lower than that in AD samples at ∼60 hours (*p*<0.001 for all isoforms; Fig. 5). Blank controls (tau isoform substrate only and no tissue seeds were added) did not show any seeding activities (data not shown). All six recombinant wild-type human tau isoforms exhibited nearly 100% specificity while sensitivity varied and ranged from 60-100% among different isoforms (Table 1).

**Table 1.** Demorgraphic characteristics of AD and non-AD patients and RT-QuIC assay with six tau isoforms.

| No. Case | Age | Sex | PMI | Neuropathological Diagnosis | RT-QuIC |  |  |  |  |  |
| --- | --- | --- | --- | --- | --- | --- | --- | --- | --- | --- |
|  |  |  |  |  | 0N3R | 1N3R | 2N3R | 0N4R | 1N4R | 2N4R |
| 1 | 17 | M | 27 | Vasculitis; Microhemorrhages | Neg | Neg | Neg | Neg | Neg | Neg |
| 2 | 35 | F | 19 | Metaoblic astrocytosis | Neg | Neg | Neg |  |  | Neg |
| 3 | 65 | M | 3 | Cerebellar degeneration; Seizure | Neg | Neg | Neg | Neg |  |  |
| 4 | 70 | M | 11 | No pathological diagnosis | Neg | Neg |  |  | Neg | Neg |
| 5 | 71 | M | 21 | A0, B1, C0 | Neg | Neg |  |  | Neg | Neg |
| 6 | 74 | M | 16 | Neocortical amyloid and neurofibrillary degeneration; Braak I/II | Neg | Neg | Neg |  |  | Neg |
| 7 | 74 | F | 6 | Frontal and temporal atrophy; arterial aneurisms, infarct | Neg | Neg |  | Neg |  |  |
| 8 | 75 | M | 16 | Argyrophilic grain disease; A1, B1, C0 |  | Neg | Neg |  |  | Neg |
| 9 | 79 | M | 17 | Microscopic infarcts; occasional NFT and amyloid plaques | Neg | Neg | Neg |  |  | Neg |
| 10 | 55 | F | 4 | AD; A3, B3, C3 | fNeg | fNeg | fNeg | Pos | Pos |  |
| 11 | 57 | M | 16 | AD; Braak VI |  |  |  | Pos | Pos | Pos |
| 12 | 63 | F | 18 | AD; A3, B3, C3 |  | Pos | Pos |  |  | Pos |
| 13 | 64 | M | 3 | AD; Braak V/VI | Pos |  |  | Pos | Pos | Pos |
| 14 | 65 | M | 38 | AD; A3, B3, C2 | fNeg | fNeg | fNeg | Pos | Pos |  |
| 15 | 67 | M | 3 | AD/LBD; many NFT, minimal amyloid plaques | Pos | Pos | Pos | Pos |  | Pos |
| 16 | 68 | M | 24 | AD/LBD |  | fNeg | Pos |  |  |  |
| 17 | 72 | M | 19 | AD/LBD; A3, B2, C2 |  | Pos | Pos |  |  | Pos |
| 18 | 74 | F | 10 | AD/LBD |  |  |  | Pos | Pos | Pos |
| 19 | 74 | M | 3 | AD |  |  |  | Pos | Pos |  |
| 20 | 74 | M | 25 | AD; Braak IV-VI |  |  |  | Pos | Pos |  |
| 21 | 78 | M | 4 | AD; Braak VI | Pos | Pos | Pos |  |  |  |
| 22 | 79 | M | 2.5 | AD/LBD | fNeg | Pos | fNeg |  |  | Pos |
| 23 | 87 | F | 3 | AD; A3, B3, C3 | Pos | Pos | Pos |  |  | fNeg |
| 24 | 87 | F | 6 | AD; A3, B2, C3 | Pos | fNeg | fNeg |  |  | Pos |
| 26 | 90 | F | 6 | AD; A3, B3, C3 |  |  | Pos |  |  | fNeg |
| 27 | 69 | M | 6 | AD; A2, B2, C3 | Pos | Pos | Pos |  |  | Pos |

Abbreviations: Braak stages are defined as in the literature (Braak & Braak, 1991). A scores (A1, A2, A3) designate different Thal phases for amyloid plaques by immunohistochemistry. B scores denote Braak stage for neurofibrillary degeneration. C scores are CERAD scores for density of neocortical neurite plaque. Neg – negative detection; Pos – positive detection; fNeg – False negative detection. LBD – Lewy body dementia.

**Figure 4.**
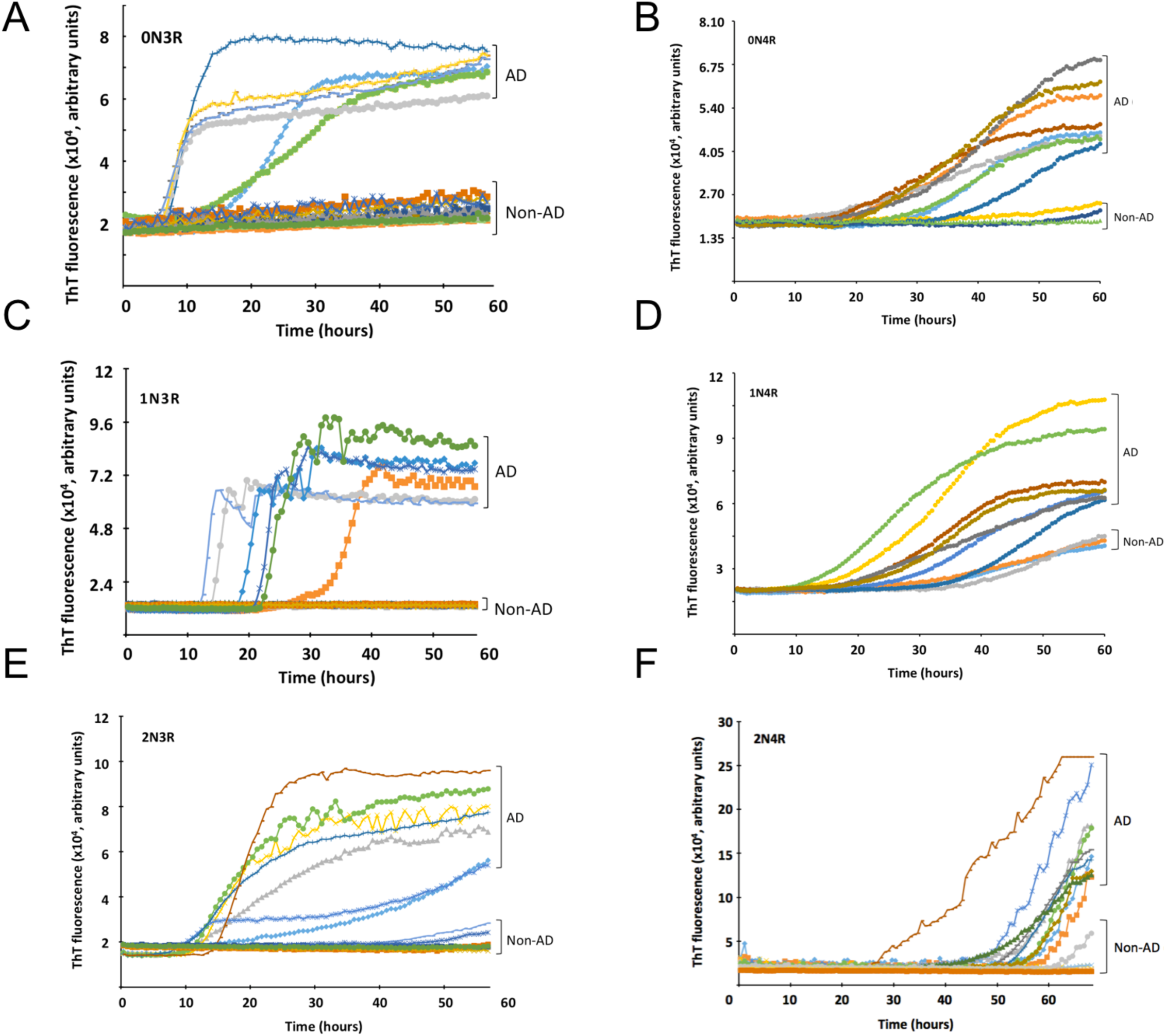
RT-QuIC analysis of tau isoform seeding with brain tissues of post-mortem AD and non-AD control subjects. Tau seeding activity of AD brains was examined by RT-QuIC assays with all six individual recombinant tau isoforms as specified. Each colored trace represented a brain homogenate sample from an individual subject.

**Figure 5.**
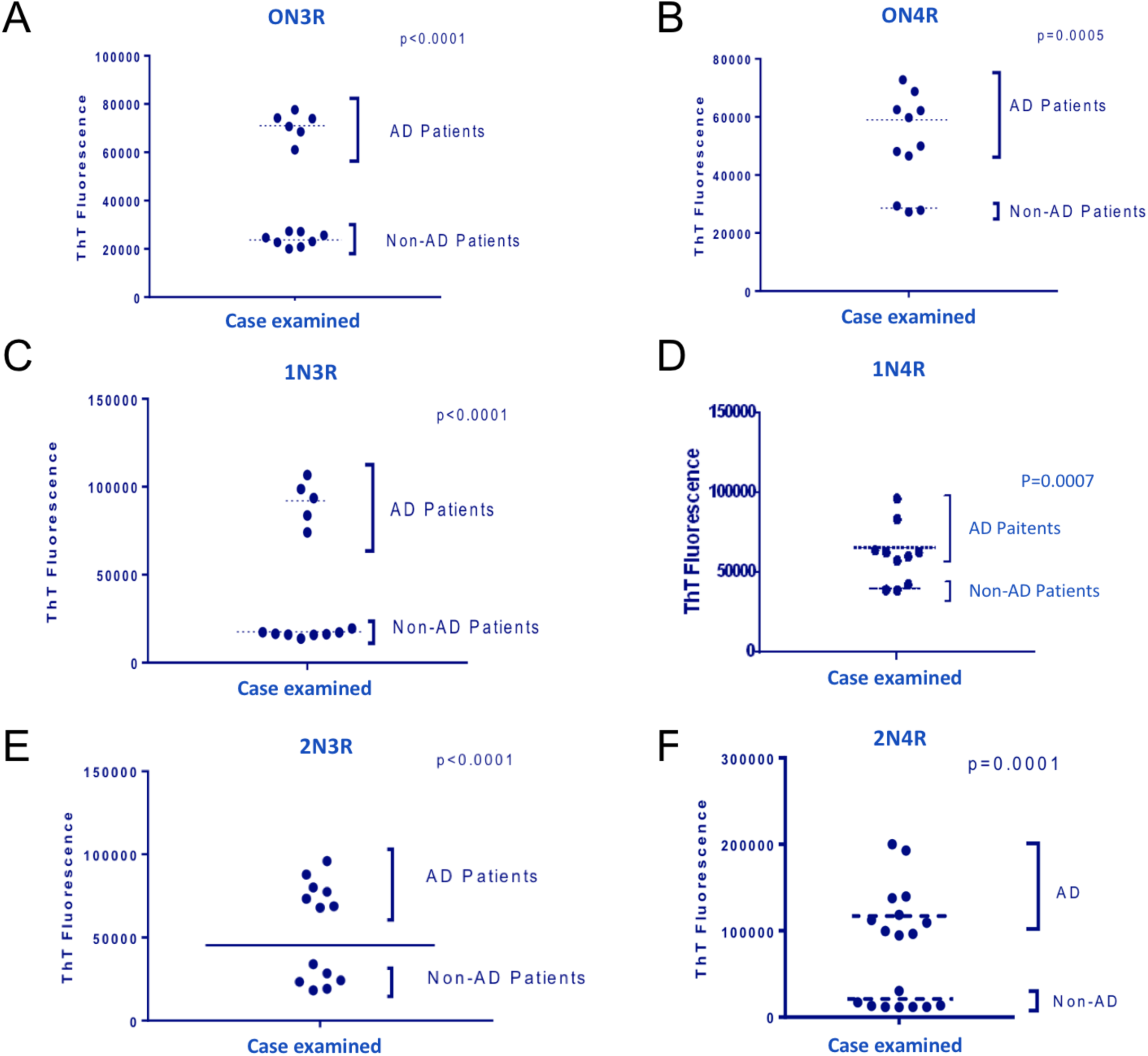
Statistical evaluation of RT-QuIC assays of tau isoform seeding with brain tissues of post-mortem AD and non-AD control subjects as described in Figure 4. GraphPad Prism 6.0 software was used for analyses. Statistical significance p-values are specified.

### Characterization of PHF-tau extracted from postmortem AD brains

Following Sarkosyl extraction-based protocols (Goedert et al, 1992; Lee et al, 1999; Guo et al, 2016), we purified enriched PHF-tau (AD tau) from postmortem AD brains. Enriched PHF-tau was clustered between 45-70 kD in migration in SDS-PAGE gel (Fig. 6A). These isoforms were confirmed using Western blotting (Fig. 6B) with an anti-tau monoclonal antibody (HT7) that targets the conserved PPGQK sequence (corresponding to residues 159-163 in 2N4R). Our Western blotting analysis showed at least three closely clustered bands (Fig. 6B), in good agreement with PHF-tau preparations described in the literature (Goedert et al, 1992; Guo et al, 2016). Additional TEM characterization of the native tissues showed abundant circular aggregates and short filament species of assembled tau isoforms (Fig. 6C).

In an effort to identify tau isoforms, we performed mass spectrometric analyses of AD brain extracted PHF-tau via a standard bottom-up analysis of released tryptic peptides. Results provided unambiguous identification of peptides found in all six isoforms although one cannot provide relative ratios for all six without using labeled standards as there are no prototypic peptides for any individual isoform (Fig. 7 and Table 2). In addition, a search that included dynamic phosphorylation of serine, threonine, or tyrosine yielded identification of doubly and triply phosphorylated peptides of HLSNVSSTGSIDMVDSPQLATLADEVSASLAK, a peptide located near the C-terminus of all six isoforms. While unambiguous evidence for the placement of all phosphates could not be accomplished, we were able to determine the percent chances of occupancy of phosphorylated sites at the C-terminus (Fig. 7). Specifically, significant phosphorylation occupancies were observed for S409, S413, T414, S416, and S422 (based on 2N4R numbering; equivalent to S3, S7, T8, S10, and S16 in the boxed insert in Fig. 7). Our findings are consistent with reported hyperphosphorylation of tau in PHF with clustered sites at the C-terminus (Morishima-Kawashima et al, 1995; Hanger et al, 1998; Hanger et al, 2007).

**Table 2.**
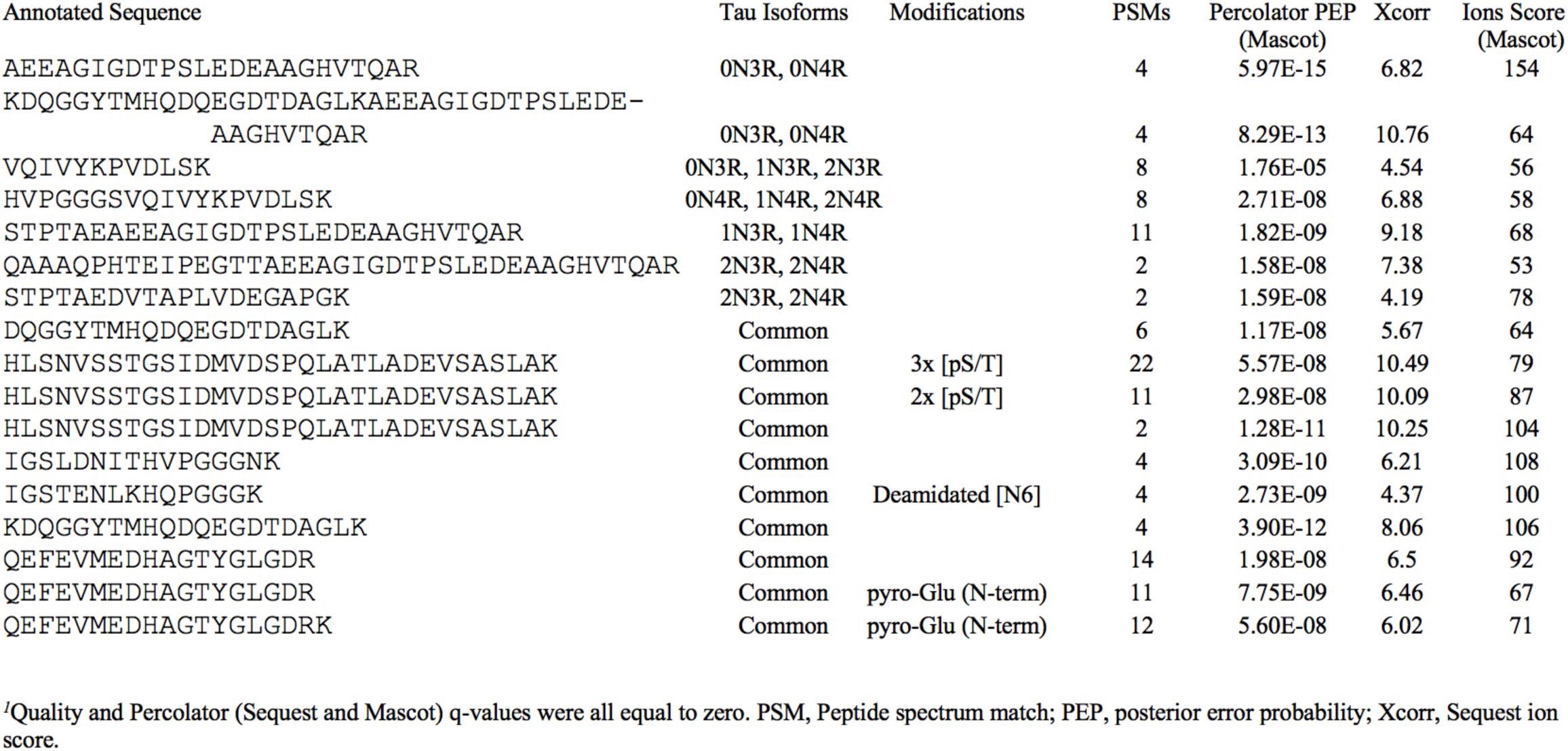
High Scoring Tryptic Tau Peptides Isolated From Alzheimer Patient Brain Tissue.^1^.

**Figure 6.**
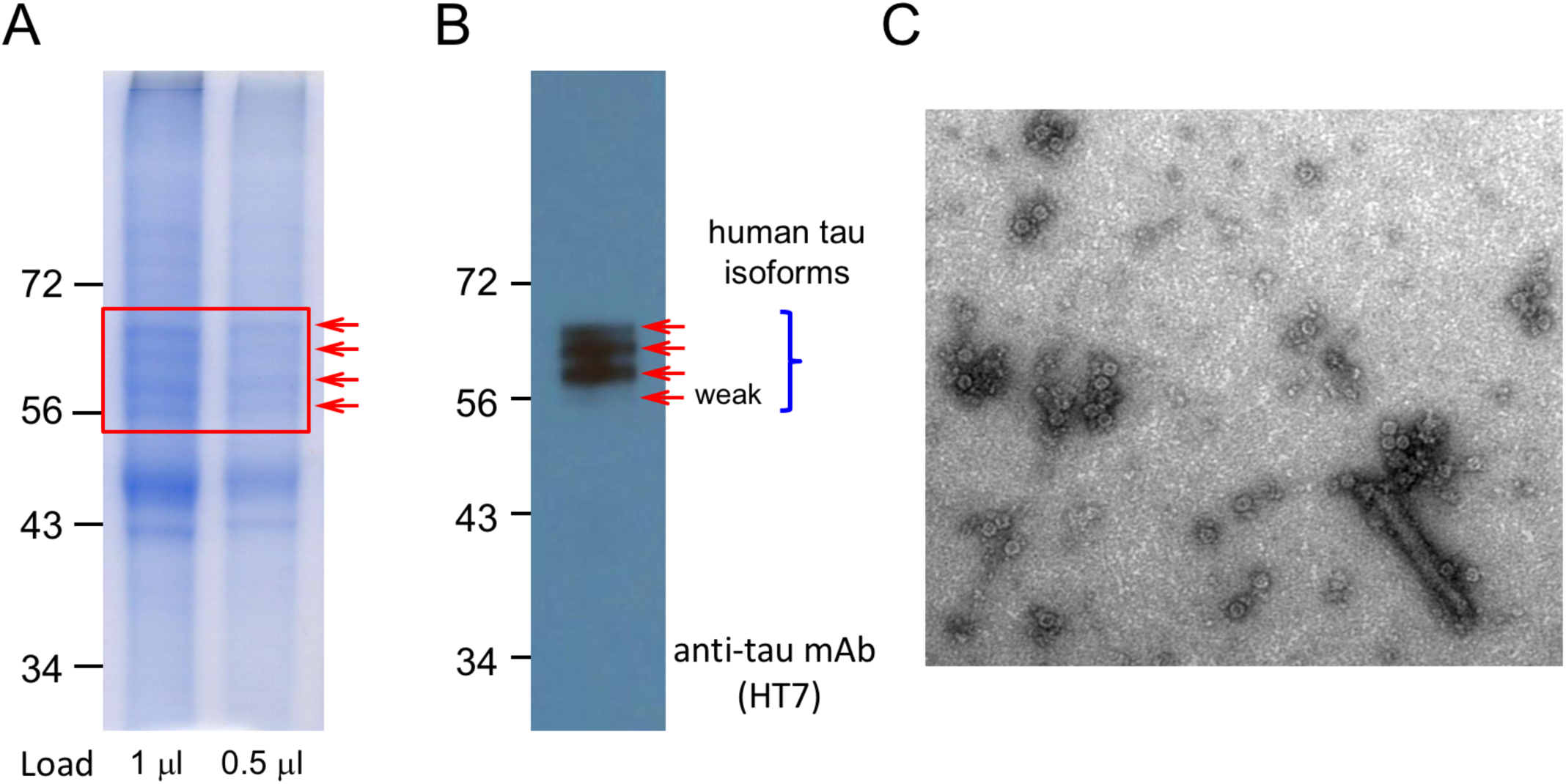
Human tau isoforms extracted from post-mortem AD brains. (A) SDS-PAGE analysis showed authentic human tau isoforms enriched from Sarkosyl extraction and differential ultracentrifugation. (B) Western blot analysis identified human tau isoforms from AD brain extracted human tau isoform preparations. Anti-tau mAb (HT7; ThermoFisher Scientific, Waltham, MA) was used for human tau isoforms detection. (C) TEM images of human tau isoform preparations extracted from AD brains. Abundant circular human tau isoform species and occasional short tau filament species were visualized after staining with uranyl acetate.

**Figure 7.**
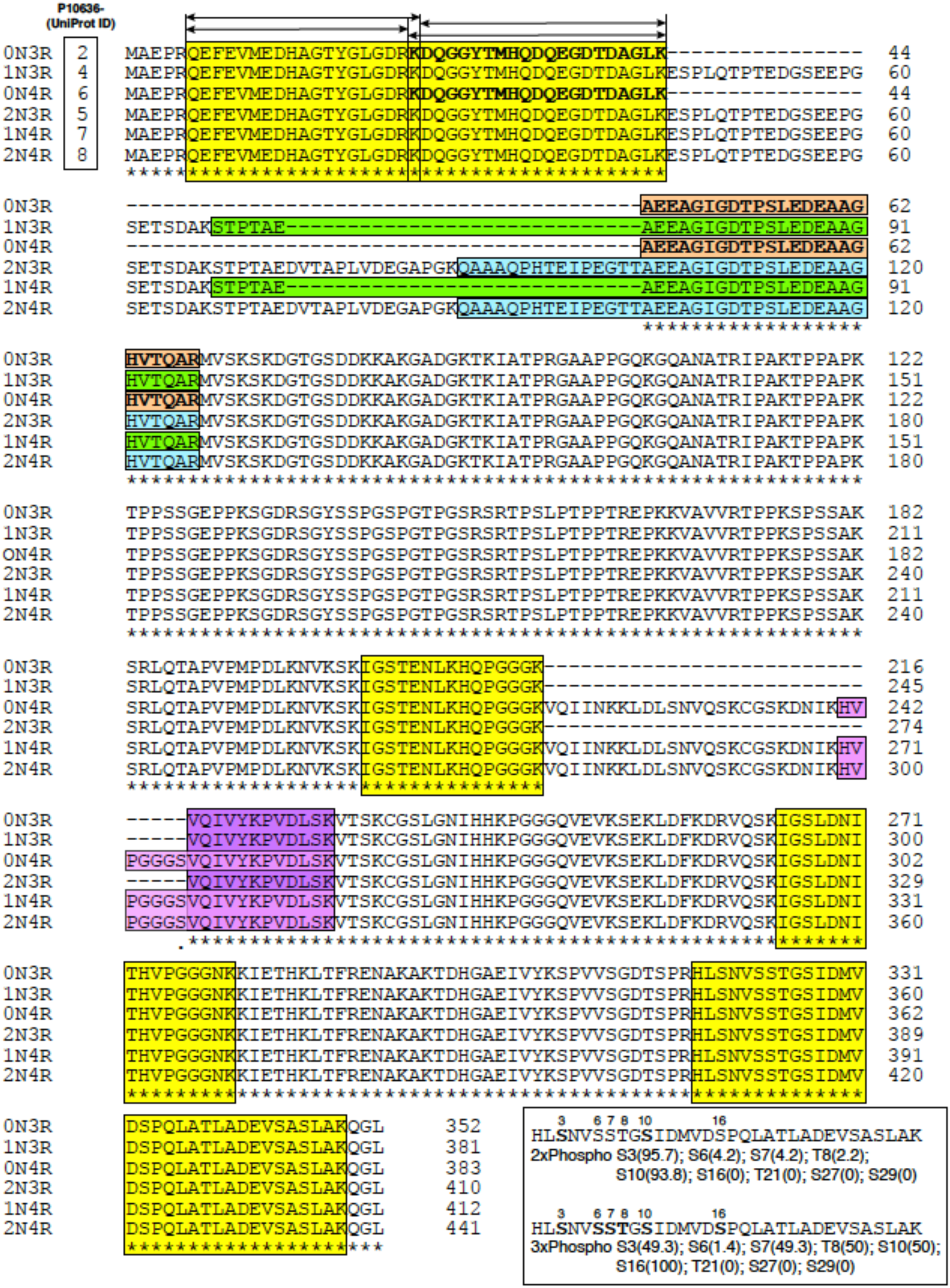
Mass spectrometric identification of human tau isoforms and phosphorylation sites of AD tau sample extracted from postmortem AD patient brain tissues. Results are summarized in the format of CLUSTAL O (1.2.4) multiple sequence alignment. Fully conserved amino acids are marked as asterisks below the sequences. Yellow boxes indicate common tryptic peptides detected. Other colors (gold, green, cyan, light and dark purple) denote peptides for specific isoforms. Same color are used for common peptide fragment detected from different isoforms. The C-terminal common peptide is multiply phosphorylated at 2 or 3 sites with percent chances of occupancy shown as determined by Proteome Discoverer (boxed insert). See Table 2 for more details.

## Discussion

Applying both concepts and assays derived from prion diseases, we were able to successfully amplify misfolded tau seeds from AD brains using various individual full-length tau isoform as substrate. To our knowledge, no studies have been reported to selectively detect misfolded tau seeds from AD brains using recombinant full-length tau isoforms as substrates. Our study complements a recent investigation describing ultrasensitive detection of tau aggregate conformer of AD with a mixture of two significantly shortened tau fragments as dual substrates (Kraus et al, 2019). The successful seeding activities of AD brain homogenates with individual tau isoforms as a substrate further validated the predicted presence of all six misfolded tau isoforms in AD brains (Goedert et al, 1992; Lee et al, 1999). Non-AD brain samples yielded minimal or significantly lower levels of fluorescent signals for aggregation. Our AD tau isoform-based RT-QuIC assays provides a promising platform for rapid diagnosis of AD and other tauopathies for future development.

Past biochemical studies and more recent cryo-EM structures of tau isoforms from AD and Pick’s diseases (PiD) suggested critical roles of the MT binding repeats in their aggregation (von Bergen et al, 2000; von Bergen et al, 2006; Fitzpatrick et al, 2017; Falcon et al, 2018). Furthermore, two minimal hexapeptide motifs at the beginning of the second and the third internal repeats were identified as two potential nucleation core sequences (von Bergen et al, 2000; Li & Lee, 2006; Seidler et al, 2018). Our systematic truncation mutagenesis of amino-terminal and carboxy-terminal mutants further suggested that at least one of the R2 or the R3 repeat is required to induce tau aggregation. Our data showed that the VQIINK/VQIVYK hexapeptide sequences (PHF6 and PHF6* sequences), but not shared VQI tripeptide sequence, are sufficient to induce tau proteins for aggregation. These data are consistent with not only the results of past proline scanning mutagenesis of PHF6 sequence (von Bergen et al, 2000), but also the insights from the cryo-EM structures of the protofilament core of the Alzheimer and Pick tau folds (Fitzpatrick et al, 2017; Falcon et al, 2018), which suggest hydrophobic forces from the entire hexapeptide sequences are necessary to drive the aggregation nucleation.

ThT fluorescence assays and orthogonal pelleting assays revealed intriguing differential aggregation kinetics of the 3R and 4R isoforms. As the differences between 0N3R/0N4R, 1N3R/1N4R, and 2N3R/2N4R are due to the presence or absence of the R2 repeat, the presence of the R2 repeat, while itself a key aggregation core element, surprisingly slows down the aggregation of the resulting isoforms. It is unknown presently why inclusion of a key aggregation element to the 3R tau isoforms will yield corresponding 4R isoforms with slower aggregation kinetics. The fold increase of t_1/2_ is also sequentially reduced from 0N3R/0N4R (12.2 fold) to 1N3R/1N4R (3.9 fold), and to 2N3R/2N4R (1.9 fold). We hypothesize that amino-terminal N1 and N2 segments play some novel roles in tau aggregation partly because the differences in t_1/2_ fold of changes may be assigned to the presence or absence of the N1 and N2 segments of the isoforms evaluated. It will be interesting to investigate the functions of the N1 and N2 segments in tau isoform aggregation and neurotoxicity in future studies.

Past studies have demonstrated that all six tau isoforms are present in tau PHFs from the brains of AD patients (Goedert et al, 1992; Lee et al, 1999). Normal adult humans express approximately equal levels of 3R-tau and 4R-tau; however, select sporadic tauopathies exhibit an imbalance of 4R-tau isoform deposition within neurofibrillary tangles (NFTs) (Arai et al, 2001; de Silva et al, 2003). Most of such semi-quantitative estimates were based on immunodetection of tau isoforms (Goedert et al, 1992; Lee et al, 1999). Yet detailed quantitative measurements of human tau isoforms within NFTs such as PHF-tau are currently not available. Our mass spectrometric analysis of PHF-tau and recent literature (Barthelemy et al, 2019) demonstrated that it is possible to perform target quantifications in normal and diseased samples if isotope-labeled peptide standards with known quantities are spiked in the testing samples and used as external standards. Such quantification may help to addresses important unresolved questions regarding 3R/4R ratio in normal and diseased CNS samples, and how imbalance of 3R/4R ratios among a combination of tau isoforms affects their aggregation and contribute to AD pathogenesis. Recent development in new technology such as ion mobility-mass spectrometry (Young et al, 2014; Young et al, 2015a, 2015b) may also provide powerful new avenues to define protein aggregate intermediates in functionally relevant tau isoform strains.

Multiple cell-based and *in vivo* mice based tau-seeding experiments provided strong evidence that tau protein has essential characteristics of a prion (Guo et al, 2011; Sanders et al, 2014; Guo et al, 2016; Dujardin et al, 2018; Sharma et al, 2018). Prion-like propagation of tau aggregates may therefor underlie the disease pathogenesis and progression of neurodegenerative tauopathies. Recent groundbreaking cryo-EM structures of tau filaments from patients with AD and Pick’s disease patients further provided atomic evidence of different molecular conformers for two distinct neurodegenerative tauopathies (Fitzpatrick et al, 2017; Falcon et al, 2018). Utilizing concepts and techniques such as RT-QuIC originally used in prion disease research, we demonstrated “prion-like” tau isoform seeding activities in diseased AD brains. Despite the progress in our understanding of how tau isoforms aggregate and propagate, many important questions remain to be addressed. For example, does tau isoform aggregation kinetics play any roles in disease progression? How do different tau isoforms, with and without post-translational modifications, interact with each other? What are the quantitative tau isoform levels in normal and diseases AD brains? Are there specific tau isoforms or isoform ratios *in vivo* that distinguish AD from normal subjects and other tauopathies? What are the molecular mechanisms that differentiate the rapid progressive and slow progressive AD diseases (Schmidt et al, 2011; Drummond et al, 2017)? New conceptual and technological advances in the field will facilitate to address these important questions.

## Materials and Methods

### Plasmid Constructs

Expression vectors for his-tagged versions of all six wild-type human tau isoforms were kindly provided by the late Dr. Lester “Skip” Binder and Dr. Nicolas Kanaan of Michigan State University. We amplified the N-terminal and C-terminal sequences by PCR and cloned into the same expression vector using Nde I and Xho I restriction sites. All the truncation mutants were designed with a his_6_-tag at their carboxy-termini to facilitate protein purification.

### Brain Tissue Collection, Handling and Analysis

The autopsy brain samples were collected and diagnosed neuropathologically at the Case Western Reserve University Alzheimer’s Disease Research Center and Case Human Tissue Procurement Facility, and National Prion Disease Pathology Surveillance Center. We have collected postmortem patient brain tissue samples from 18 AD patients (mean year of age 67.9 ± 11.2) and 9 non-AD controls (mean year of age 62.2 ± 19.8). The mean postmortem interval was 12.7 ± 10 hours. Demographics and Braak staging of AD-tau are described in details in Table 1 (Cohen et al, 2015) followed guidelines from the National Institute on Aging-Alzheimer’s Association workgroups (McKhann et al, 2011). All protocols currently in place were to support and facilitate the research in the context of IRB and HIPAA regulations, by articulating and implementing criteria allowing the acquisition, storage and distribution for research use of such data and specimens with or without linkage to personal health identifiers. The use of postmortem brain tissues for research was approved by the Case Western Reserve University’s Institutional Review Board with informed consent from patients or their families. Brain tissues were homogenized at 10% (w/v) in 1 × PBS with mini Beads Beater. The brain homogenates were further subjected to centrifugation at 500 × g for 10 min at 4 °C and the supernatants were collected and stored at −80 °C for future RT-QuIC assays as seeds (see below).

### Recombinant Human Tau Isoform Expression and Purification

Plasmids encoding human tau isoforms or truncation mutants were transformed into BL21-DE3 *E. coli* cells. Overnight starter cultures of recombinant human tau isoforms or variants were prepared using 100 mL of LB broth with 100 μg/mL ampicillin inoculated with recombinant competent cells from transformation procedure and grown at 37°C, shaking at 180 RPM. Four flasks containing 1 liter of LB broth were inoculated with 20 mL of starter culture and 100 mg/mL ampicillin. Cultures were incubated at 37°C, shaking until OD_600_ reached between 0.5-0.8. Tau expression was induced using 1 mM IPTG and continued to grow for an additional 4 hours. BL21-DE3 cells containing expressed tau were pelleted via centrifugation at 8,500 RPM for 20 minutes at 20 °C. Pelleted cells were resuspended in 50 mM NaH_2_PO_4_, pH 8 and 300 mM NaCl (sonication lysis buffer) at a concentration of 20 mL/L of culture preparation and sonicated at 60% power in ten 30 second intervals over 10 minutes. Cell lysates were centrifuged at 8,500 RPM for 20 minutes (an extra boiling step at 90 °C for 20 minutes was added for all tau isoforms before centrifugation; Barghorn et al, 2005). Supernatant containing protein was applied to Ni-NTA column equilibrated with sonication lysis buffer. The columns were washed with 30-50 times of bed volumes of column buffer followed by washing buffer (50 mM NaH_2_PO_4_, pH 8.0, 300 mM NaCl, and 30 mM imidazole). Recombinant protein was then eluted using elution buffer (50 mM NaH_2_PO_4_, pH 8.0, 300 mM NaCl, and 200 mM imidazole). Fractions were tested for protein concentration using 5 μL of protein sample mixed with 10 μL Coommassie Protein Assay reagent (Thermo Scientific). Pooled fractions were concentrated to 4 mL using 10 kD molecular weight cut-off spin columns (Millipore) and filtered using 0.22 μm low-binding Durapore PVDF membrane filters (Millipore). Tau protein was further purified by FPLC using size exclusion Superdex75 and Superdex200 columns (GE Healthcare) in 1X PNE buffer (25 mM PIPES pH 7.0, 150 mM NaCl and 1 mM EDTA).

### Purification of Tau PHFs from AD Brains

PHF-tau purification follows established protocols (Goedert et al, 1992; Lee et al, 1999; Guo et al, 2016). Briefly, brain tissues from sporadic AD patients with abundant tau pathology (Braak stage VI) qualified for AD were used in this study. All cases used were histologically confirmed. For each purification, 10-12 g of frontal cortical gray matter was homogenized using a homogenizer in multiple volumes (v/w) of high-salt buffer (10 mM Tris-HCl, pH 7.4, 0.8 M NaCl, 1 mM EDTA, and 2 mM dithiothreitol (DTT), with protease inhibitor cocktail, with 0.1% Sarkosyl and 10% sucrose added and centrifuged at 10,000 *g* for 10 min at 4°C. Pellets were reextracted once or twice using the same buffer conditions as the starting materials, and the supernatants from all two to three initial extractions were filtered and pooled. Additional Sarkosyl was added to the pooled low-speed supernatant to reach 1%. After one-hour nutation at room temperature, samples were centrifuged again at 300,000 *g* for 60 min at 4°C. The resulted 1% Sarkosyl-insoluble pellets, which contain pathological tau, were washed once in PBS and then resuspended in PBS (∼100 µl/g gray matter) by passing through 27-G 0.5-inch needles. The resuspended Sarkosyl-insoluble pellets were further purified by a brief sonication (20 pulses at ∼0.5 s/pulse) followed by centrifugation at 100,000 *g* for 30 min at 4°C, whereby the majority of protein contaminants were partitioned into the supernatant, with 60–70% of tau remaining in the pellet fraction. The pellets were resuspended in PBS at one fifth to one half of the pre-centrifugation volume, sonicated with 20–60 short pulses (∼0.5 s/pulse), and spun at 10,000 *g* for 30 min at 4°C to remove large debris. The final supernatants, which contained enriched AD PHFs, were used in the study and referred to as AD-tau. The final supernatant fraction was further analyzed by SDS-PAGE, Western blotting, transmission electron microscopy, BCA assay, and mass spectrometry.

### Thioflavin-T Fluorescence Aggregation Kinetic Analysis

Fluorescence experiments were performed using a SpectraMax M5 plate reader (Molecular Devices, Sunnyvale, CA). All kinetic reads were taken at 37 °C in non-binding all black clear bottom Greiner 96-well plates covered with optically clear films and stirred for 10 seconds prior to each reading. ThT fluorescence was measured at 444 nm and 491 nm as excitation and emission wavelengths. Each kinetic assay consisted of final concentrations of 30 μM tau protein and 10 μM ThT. The amount of time required to reach half maximum ThT intensity (t_1/2_) and inhibition constant (IC_50_) values for dose response curves were estimated by multiparameter logistic nonlinear regression analysis. The transition from the lag-phase to the growth phase was estimated when the first measurable ThT fluorescence/time (slope) value exceeded ≥ 5 fold of the previous measured slope (i.e. where the quantum leap in ThT RFU is first noticeable).

### Sarkosyl-insoluble Tau Pelleting Assay

Purified recombinant tau (10 μM) and heparin (100 μg/ml) were incubated at 37 °C for 48 h in 30 mM Tris-HCl, pH 7.5, containing 2 mM DTT. Aggregated tau was assayed on the basis of 1% Sarkosyl insolubility as described below. Aliquots (10 μl) of assembly mixtures were removed and added to 50 μl of 30 mM Tris-HCl, pH 7.5, containing 1% Sarkosyl, and the mixture was left for 30 min. The mixture was then spun at 150,000 × *g* for 20 min. The supernatant (Sarkosyl-soluble tau) was removed, and the pellet (Sarkosyl-insoluble tau) was resuspended in 20 μl of SDS sample buffer containing 5% 2-mercaptoethanol and subjected to SDS-PAGE. Aggregated tau isoforms were visualized after staining of the gel with Coomassie Brilliant Blue.

### Transmission Electron Microscopy (TEM) Analysis

TEM images were collected as previously described (Velander et al, 2016; Wu et al, 2017). Briefly, 30 μM human tau isoforms were incubated in 20 mM Tris-HCl, pH 7.4 for 10-12 days at 37 °C. Prior to imaging, 2 μL of sample were blotted on a 200 mesh formvar-carbon coated grid for 5 minutes, and stained with uranyl acetate (1%) for 1 minute. Both sample and stain solutions were wicked dry (sample dried before addition of stain) by filter paper. Qualitative assessments of the amount of fibrils or oligomers observed were made by taking representative images following a careful survey of each grid. At least 15-20 locations of each grid were surveyed in collecting representative images of each sample. TEM was performed on a JEOL-1400 transmission electron microscope (JOEL USA, Inc., Peabody, MA) operated at 120 kV.

### RT-QuIC Analysis

Tau RT-QuIC assays were conducted as previously described with minor modifications (Orru et al., 2017; Saijo et al., 2017; Kraus et al., 2019; Wang et al., 2019). In brief, each recombinant tau isoform was thawed at room temperature and filtered with a 100 kDa spin column filter (Millipore) to remove unwanted tau aggregates. RT-QuIC reaction mix was composed of 40 μM heparin, 10 mM HEPES, 400 mM NaCl, and 10 μM thioflavin T (ThT) and was filtered through a 0.22 µm filter before use. Each tau isoform was slowly and gently added into the RT-QuIC reaction mix at a final concentration of 20 µM. A 98 μl of RT-QuIC reaction mix containing tau isoform was loaded into each well of a 96-well plate (Nunc). The 10% brain homogenates were diluted at 1:200 in a dilution buffer containing 10 mM HEPES and 1.7% 1 × N2 supplement (Gibco) and centrifuged at 2000 g for 2 min at 4°C. A 2 µl of diluted brain homogenate was added to the 98 µl of RT-QuIC reaction mix in each well. The plate was then sealed with a plate sealing tape (Nalgene Nunc International) and incubated at 37°C in a BMG FLUOstar Omega plate reader (BMG Labtech FLUOstar Omega) with cycles of 1 min shaking (500 rpm orbital) and 1 min rest throughout the incubation time. ThT fluorescence measurements were monitored every 45 min using 450 +/-10 nm excitation and 480 +/-10 nm emission by bottom reading. Instrument gains were set at 2016. Four replicate reactions were prepared at the same dilution for each individual sample. The average fluorescence values per sample were calculated using fluorescence values from all four replicate wells. At least 2 of 4 replicate wells must cross this threshold for a sample to be considered as positive.

### Mass Spectrometric Analysis

PHF-tau extracted from neuropathologically diagnosed AD patient brain cortex tissues were tryptic digested following standard protocol (Lee et al, 1999; Guo et al, 2016). Mass spectrometric analysis utilized a Waters Synapt G2-S HDMS (Waters, Milford, MA) interfaced with an Acquity I-Class UPLC system was operated in continuum positive ion “resolution” MS mode using an HDMSE acquisition method (high-definition mass spectrometry with alternating scans utilizing low and elevated collision energies with ion mobility separation (IMS) of peptides prior to fragmentation). Both low energy (4 V and 2 V in the trap and transfer region, respectively) and elevated energy (4 V in the trap and ramped from 20 to 50 V in the transfer region) scans were 1.2 seconds each covering the m/z range of 50 to 1800. For ion mobility separation, the IMS and transfer wave velocities were 600 and 1200 m/sec, respectively. Wave height within the ion mobility cell was ramped from 10 to 40 V. For lock-mass correction, a 1.2 second low energy scan was acquired every 30 seconds of a 100 fmol/μL [Glu1]-fibrinopeptide B (Waters, Milford, MA) solution (50:50 acetonitrile:water supplemented with 0.1% (v/v) formic acid) infused at 10 μL/min introduced into the mass spectrometer via a different source also utilizing a capillary voltage of 3 kV.

### Statistical Analysis

All data are presented as the mean ± S.E.M and the differences were analyzed with unpaired Student’s *t* test as implemented within GraphPad Prism software (version 6.0). *p* values < 0.05 were considered significant. Tau RT-QuIC data were plotted and analyzed using GraphPad Prism as well. The seeding activity was considered to be tau positive if the ThT fluorescence exceeded a threshold reading. The threshold was defined as the average of mean baseline readings over the 30-60 h time window in the tau RT-QuIC traces for the non-tauopathy samples in the given experiments plus 20 times the standard deviation of those readings.

## Acknowledgments

We thank Case Western Reserve University Alzheimer’s Disease Research Center (P30 AG062428), Case Human Tissue Procurement Facility, and National Prion Disease Pathology Surveillance Center for AD and non-AD brain tissues access. We thank Kathy Lowe at Virginia-Maryland Regional College of Veterinary Medicine for her excellent technical assistance in using transmission electron microscope. We thank Virginia Tech Center for Drug Discovery for instrument access.

## Availability of data and materials

The datasets generated during this study are available from the corresponding authors upon request.

## Author contributions

LW and ZW performed experiments, acquired and analyzed the data. SL, DTD, SSM, MM, FH, CT assisted with the experiments. RFH designed and analyzed the mass spectrometric experiments and WKR performed and analyzed the experiments. JL participated in the experiments and provided inputs in experimental design. XZ and SS contributed key reagents and clinical diagnosis data. GSB contributed key reagents and provided inputs in experimental design. WZ collected brain tissues, designed, acquired and analyzed the data, and prepared the manuscript. BX conceived, designed the experiments, analyzed the data, and prepared the manuscript. All the authors approved the final manuscript.

## Ethics approval and consent to participate

All procedures were performed with informed consent in accordance with state laws and institutional review board guidelines of Case Western Reserve University.

