## Supplemental Table 1 for "Human Tau Isoform Aggregation and Selective Detection of Misfolded Tau from Post-Mortem Alzheimer’s Disease Brains"

**Supplemental Information**

**Table S1**. PCR oligo primers for cloning of all the tau truncation mutants are listed. Please note all constructs except C7 were cloned using 2N4R tau as the template. 2N3R tau was used as the template for generating the C7 construct.

| Plasmid construct | Forward Primer | Reverse Primer |
|---|---|---|
| N1 | CATTCATATGGAATCTCCCCTGCAG | TTAACTCGAGTCAATGATGATGATGATGATGCAAA CCCTGCTTGGCCAGG |
| N2 | CATTCATATGGTGACAGCACCCTTA | TTAACTCGAGTCAATGATGATGATGATGATGCAAA CCCTGCTTGGCCAGG |
| N3 | CATTCATATGGAAGAAGCAGGCATT | TTAACTCGAGTCAATGATGATGATGATGATGCAAA CCCTGCTTGGCCAGG |
| N4 | AATTCATATGCAGACAGCCCCCGTGCCCAT | TTAACTCGAGTCAATGATGATGATGATGATGCAAAC CCTGCTTGGCCAGG |
| N5 | TTAACATATGGTGCAGATAATTAATAAG | TTAACTCGAGTCAATGATGATGATGATGATGCAAAC CCTGCTTGGCCAGG |
| N6 | TTAACATATGGTGCAAATAGTCTAC | TTAACTCGAGTCAATGATGATGATGATGATGCAAAC CCTGCTTGGCCAGG |
| N7 | TTAACATATGGTGGAAGTAAAATC | TTAACTCGAGTCAATGATGATGATGATGATGCAAAC CCTGCTTGGCCAGG |
| N8 | TTAACATATGAAAAAGATTGAAAC | TTAACTCGAGTCAATGATGATGATGATGATGCAAAC CCTGCTTGGCCAGG |
| C1 | CATTCATATGGCTGAGCCCCGCCAG | TTAACTCGAGTCAATGATGATGATGATGATGATTT CCTCCGCCAGGGACGT |
| C2 | CATTCATATGGCTGAGCCCCGCCAG | TTAACTCGAGTCAATGATGATGATGATGATGCTGG CCACCTCCTGGTTTAT |
| C3 | CATTCATATGGCTGAGCCCCGCCAG | TTAACTCGAGTCAATGATGATGATGATGATGACTG CCGCCTCCCGGGACGT |
| C4 | CATTCATATGGCTGAGCCCCGCCAG | TTAACTCGAGTCAATGATGATGATGATGATGCTTC CCGCCTCCCGGCTGGT |
| C5 | CATTCATATGGCTGAGCCCCGCCAG | TTAACTCGAGTCAATGATGATGATGATGATGCAGGC GGCTCTTGGCGGAAG |
| C6 | CATTCATATGGCTGAGCCCCGCCAG | TTAACTCGAGTCAATGATGATGATGATGATGCTTA TTAATTATCTGCACCTTC |
| C7 | CATTCATATGGCTGAGCCCCGCCAG | TTAACTCGAGTCAATGATGATGATGATGATGTTTGT AGACTATTTGCACCTTC |
| C8 | CATTCATATGGCTGAGCCCCGCCAG | TTAACTCGAGTCAATGATGATGATGATGATGTATC TGCACCTTCCCGCCTC |
